# AAV Capsid Screening for Translational Pig Research Using a Mouse Xenograft Liver Model

**DOI:** 10.1101/2024.05.29.596409

**Authors:** Melanie Willimann, Amita Tiyaboonchai, Kei Adachi, Bin Li, Lea Waldburger, Hiroyuki Nakai, Markus Grompe, Beat Thöny

## Abstract

In gene therapy, delivery vectors are a key component for successful gene delivery and safety, based on which adeno-associated viruses (AAVs) gained popularity in particular for the liver, but also for other organs. Traditionally, rodents have been used as animal models to develop and optimize treatments, but species and organ specific tropism of AAV desire large animal models more closely related to humans for preclinical in-depth studies. Relevant AAV variants with the potential for clinical translation in liver gene therapy were previously evolved *in vivo* in a xenogeneic mouse model transplanted with human hepatocytes. Here, we selected and evaluated efficient AAV capsids using chimeric mice with a >90% xenografted pig hepatocytes. The pig is a valuable preclinical model for therapy studies due to its anatomic and immunological similarities to humans. Using a DNA-barcoded recombinant AAV library containing 47 different capsids and subsequent Illumina sequencing of barcodes in the AAV vector genome DNA and transcripts in the porcine hepatocytes, we found the AAVLK03 and AAVrh20 capsid to be the most efficient delivery vectors regarding transgene expression in porcine hepatocytes. In attempting to validate these findings with primary porcine hepatocytes, we observed capsid-specific differences in cell entry and transgene expression efficiency where the AAV2, AAVAnc80, and AAVDJ capsids showed superior efficiency to AAVLK03 and AAVrh20. This work highlights intricacies of *in vitro* testing with primary hepatocytes and the requirements for suitable pre-clinical animal models but suggests the chimeric mouse to be a valuable model to predict AAV capsids to transduce porcine hepatocytes efficiently.

## Introduction

Technologies for genetic modification are indispensable tools in research and have become promising strategies to treat genetic disorders. The field of gene therapy has progressed rapidly in the past two decades, with which safer and more efficient vectors are being developed^1^. For *in vivo* gene therapy, recombinant adeno-associated viruses (rAAVs) have emerged to be the most popular clinical delivery vectors due to their advantageous safety characteristics and high, tropism-guided transduction efficiency^1–3^. They are small, non-enveloped, self-replication incompetent viruses of the *Parvoviridae* family, depending on either adenoviruses or herpes simplex viruses for replication^4^. Encapsulated is a 4.7-kb single-stranded linear DNA genome, which is expressed episomally in host cells but in rare cases may also integrate into the chromosomes^5^. As of today, five rAAV-based gene therapies are approved by the U.S. Food and Drug Administration for *in vivo* application^6^, Luxturna (AAV2-*RPE65*), Zolgensma (AAV9-*SMN1*), Hemgenix (AAV5-*hFactorIX*), Elevidys (AAVrh74-*micro-dystrophin*), and Roctavian (AAV5-*hFactorIIX*). Many more are in clinical trials and development. Especially the liver has been shown to be a promising target organ for *in vivo* rAAV-based gene therapy to treat acquired and inherited metabolic disorders as well as clotting disorders^1, 7^ as Hemgenix does.

Laboratory mouse models have thus far provided pivotal insights into the feasibility and optimization of therapeutic approaches. However, the lack of exposure to pathogens and the significant differences in their immune system compared to humans have posed increasing bias and risks as the development of AAV-based clinical treatments progresses^8–10^. Besides immunological safety concerns, an additional layer of complexity is added by AAV’s tissue- and species-specific tropism^1, 4, 11^. Therefore, more suitable large animal models, such as non-human primates (NHP), canines, and pigs^3, 12–15^, have been utilized for preclinical studies. In particular pigs, which are known to be genetically, anatomically, physiologically, and immunologically closely related to humans, have been characterized extensively^16–18^. In many cases, they are more accessible than NHPs. Thus, they have been used as pre-clinical models to test the suitability of various AAV vectors to target a vast range of cells, including motor neurons (AAV9 and AAVhu68)^19, 20^, lung epithelial cells (AAV8, AAV2 and AAV2H22)^21, 22^, myocardial cells (AAV1 and AAV2)^23^, and photoreceptors (AAV2, AAV5, and AAV8)^24–26^. Additionally, pigs can be genetically engineered, allowing them to optimally represent specific human diseases or even be viable organ donors for humans^27, 28^.

AAVs were originally discovered as byproduct or contaminant during adenovirus production^29, 30^ and later isolated from humans^31^. Since then, various approaches have shown to successfully increase species and tissue tropism and decrease rAAV’s antigenicity to escape from pre-existing anti-AAV immunity. The most elementary method is to isolate novel AAV serotypes from different species, which until today is a viable source for highly efficient and novel AAV vectors, including AAVrh1-rh37^32^ and AAVhu1-hu67^33^ derived from rhesus macaques and humans, respectively. Upon sequencing and deciphering the AAV genome in the early 80’s, AAV capsids have successfully been engineered by capsid shuffling, short peptide insertion and bioinformatic approaches, like AAVAnc80, which often involves direct evolution^11, 34, 35^. Error prone PCR, capsid shuffling, and Cre recombination-based targeted evolution have successfully yielded optimized rAAV capsids for the liver, e.g. AAVDJ^36, 37^. Furthermore, large animal models, including pigs, are ideal hosts for directed *in vivo* evolution^38^ to develop and test new rAAV serotypes for a given tissue or organ of interest but require substantial amounts of resources. Thus, the chimeric mouse model fumarylacetoacetate hydrolase (*Fah^-/-^*), *Rag2^-/-^, Ilr2g^-/-^* NOD-(*FRGN*) mouse^39, 40^ has been used to create and screen for novel, engineered rAAV capsids highly efficient in transducing the liver^41–43^, such as AAVLK03^41^.The combination of immune deficiency and tyrosinemia type I enables xenotransplantation and repopulation of the liver with a wide variety of species. The resulting chimeric mice require less viral resources to screen or test novel liver-specific rAAV vectors.

In this study, we used chimeric *FRGN* mice repopulated with porcine hepatocytes to find suitable AAV capsids to efficiently transduce porcine hepatocytes. Intravenous administration of a DNA-barcoded AAV library^44^ into xenograft mice allowed the screening of 47 different capsids quantified by Illumina sequencing and found AAVLK03 and AAVrh20 to be the most promising rAAV capsids for porcine hepatocytes. Subsequent *in vitro* investigation provides insight into capsid-specific intracellular processing and highlights the challenges of using primary hepatocytes thus emphasizing the requirement of suitable preclinical animal models for gene therapy research.

## Results

### Strategy for ranking AAV capsids for their ability to transduce porcine hepatocytes using a chimeric mouse model

To find the most efficient AAV capsids for porcine hepatocytes among a selected capsid library, we decided to establish a model based on the *FRGN* chimeric mouse, schematically summarized in Figure 1. *FRGN* mice^39, 40^ are highly immune compromised and have tyrosinemia type I, allowing freshly isolated xenotransplanted porcine hepatocytes to engraft with minimal cell rejection and repopulate the liver upon Nitisinon (NTBC) removal. Highly (>80%) repopulated chimeric mice allowed us to screen an AAV library of 47 DNA-barcoded capsids for its most efficient AAV capsid to transduce porcine hepatocytes. Upon intravenous injection, AAV-treated chimeric mice were maintained for two weeks to allow sufficient time for intracellular processing, capsid unfolding, and subsequent DNA-barcode expression. After hepatocyte isolation, porcine hepatocytes were purified by magnetic-activated cell sorting (MACS) prior to DNA and RNA extraction followed by Illumina barcode sequencing. This approach limits the background interference of AAV capsids that transduced murine hepatocytes. Viral genome copies and corresponding mRNA copies were then quantified using next-generation sequencing (NGS) to determine the AAV capsid with the highest relative efficiency to transduce porcine hepatocytes.

**Figure 1:**
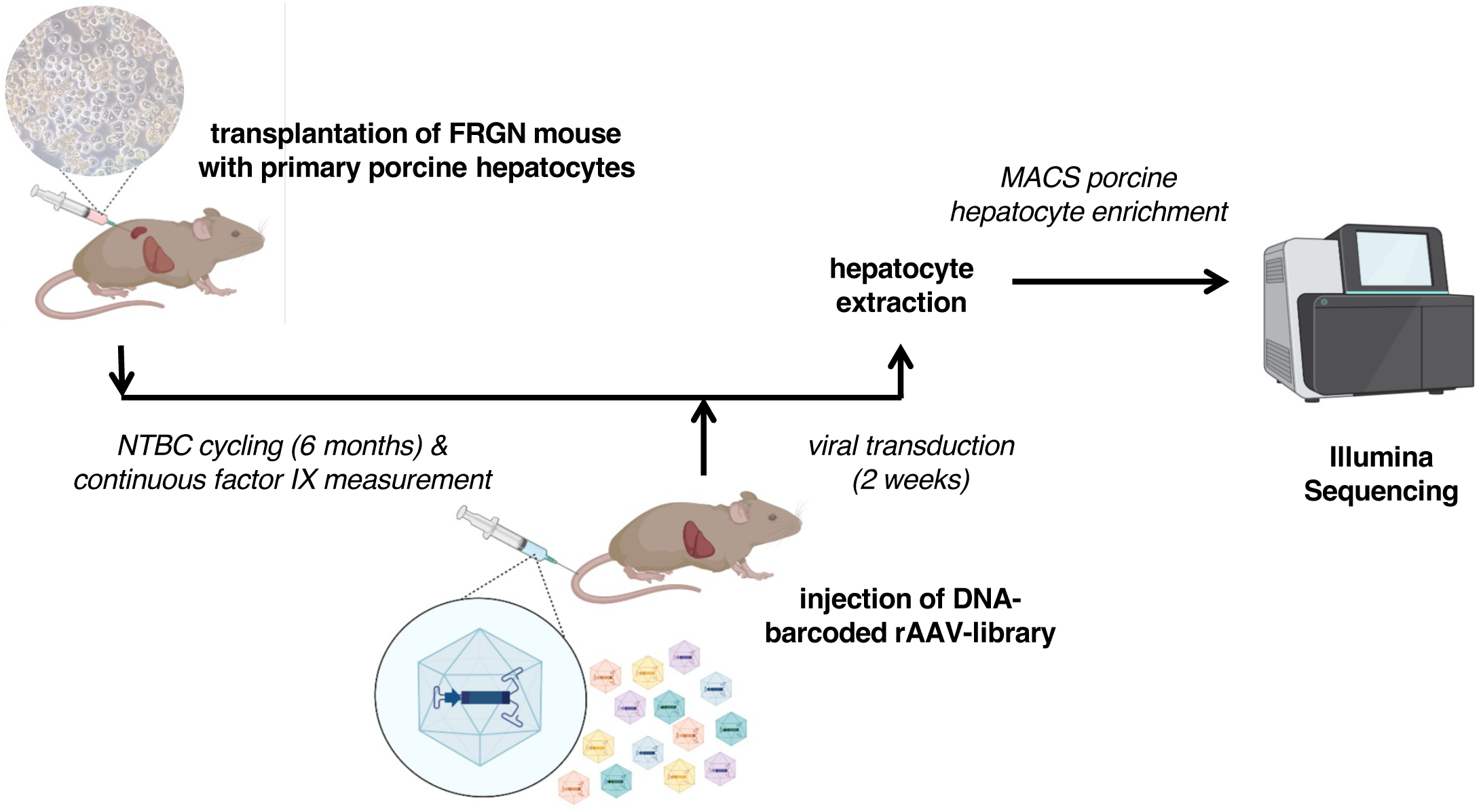
Timeline of high throughput investigation to find most efficient AAV serotype with tropism for porcine liver. 5-8 week old Fah-/-, Rag2-/-, Il2rg-/-, NOD (FRGN) mice were xenotransplanted with primary porcine hepatocytes and repopulated over the period of 5-6 months. Once xenotransplanted FRGN mice were >90% repopulated, a DNA-barcoded rAAV-library of 47 capsids was injected. 2 weeks post-injection, hepatocytes were harvested, enrichment for porcine hepatocytes using MACS, followed by DNA and RNA extraction for Illumina Sequencing, and FACS validation.

### Parameters for porcine repopulation in mice

The repopulation procedure of *FRGN* mice with primary porcine hepatocytes (1°PH) was adapted from the protocol established for robust repopulation using primary human hepatocytes. The protocol established by Azuma *et al.* included adenovirus expressing urokinase-type plasminogen activator (Ad:uPA)^39^ administration 24-48 hours prior to transplantation of human primary hepatocytes, on-off NTBC cycling, and repeated human serum albumin measurement to monitor the degree of repopulation. Various parameters were refined for xenotransplantation into *FRGN* mice with 1°PH. These refinements included skipping the pre-administration of Ad:uPA, comparing several 1°PH donors for transplantation, and monitoring the degree of repopulation by detecting porcine clotting factor IX (pFIX). Ad:uPA was administered 24 hours prior to transplantation to induce degradation of the murine hepatic extracellular matrix providing a selective advantage for the xenograft 1°PH. In total, 40 *FRGN* mice were transplanted (Supplementary Table S1). Among them, only 11% (2/18) repopulated when pre-administered Ad:uPA, whereas the repopulation success was 31% (7/22) without Ad:uPA pre-treatment. Thus, skipping Ad:uPA pre-administration was three times more successful than employing it to repopulate *FRGN* mice using porcine hepatocytes. We also compared the ability of commercial and non-commercial 1°PH to repopulate the liver, with a focus on animals that did not receive Ad:uPA. While the viability of self-isolated 1°PH was only 50-60% compared to >80% of commercially available 1°PH, the repopulation of the mouse liver was comparable, *i.e.* 31% (6/19) and 33% (1/3), respectively. Throughout NTBC cycling, pFIX secreted by the transplanted porcine hepatocytes was continuously measured to monitor the degree of repopulation. It has previously been shown that the degree of repopulation affects the experimental outcome^48^, emphasizing the importance of monitoring and confirming successful repopulation. Due to high homology of the clotting factor IX between humans and pigs, an ELISA kit established for human factor IX (hFIX) was reliably used to detect pFIX but not murine factor IX (Figure 2A). Once the pFIX levels of the xenotransplanted *FRGN* mice was >1000 ng/mL for two subsequent measurements, three mice were injected with the DNA-barcoded rAAV-library. After harvesting the hepatocytes two weeks post-injection, fluorescence-activated cell sorting (FACS) revealed that the xenograft livers were successfully repopulated for more than >90% (Figure 2B; Supplementary Table S2). Additionally, we observed that the weight of the repopulated livers accounted for 15-20% of total body weight, which is three to four times the relative liver weight compared to non-transplanted mice.

**Figure 2:**
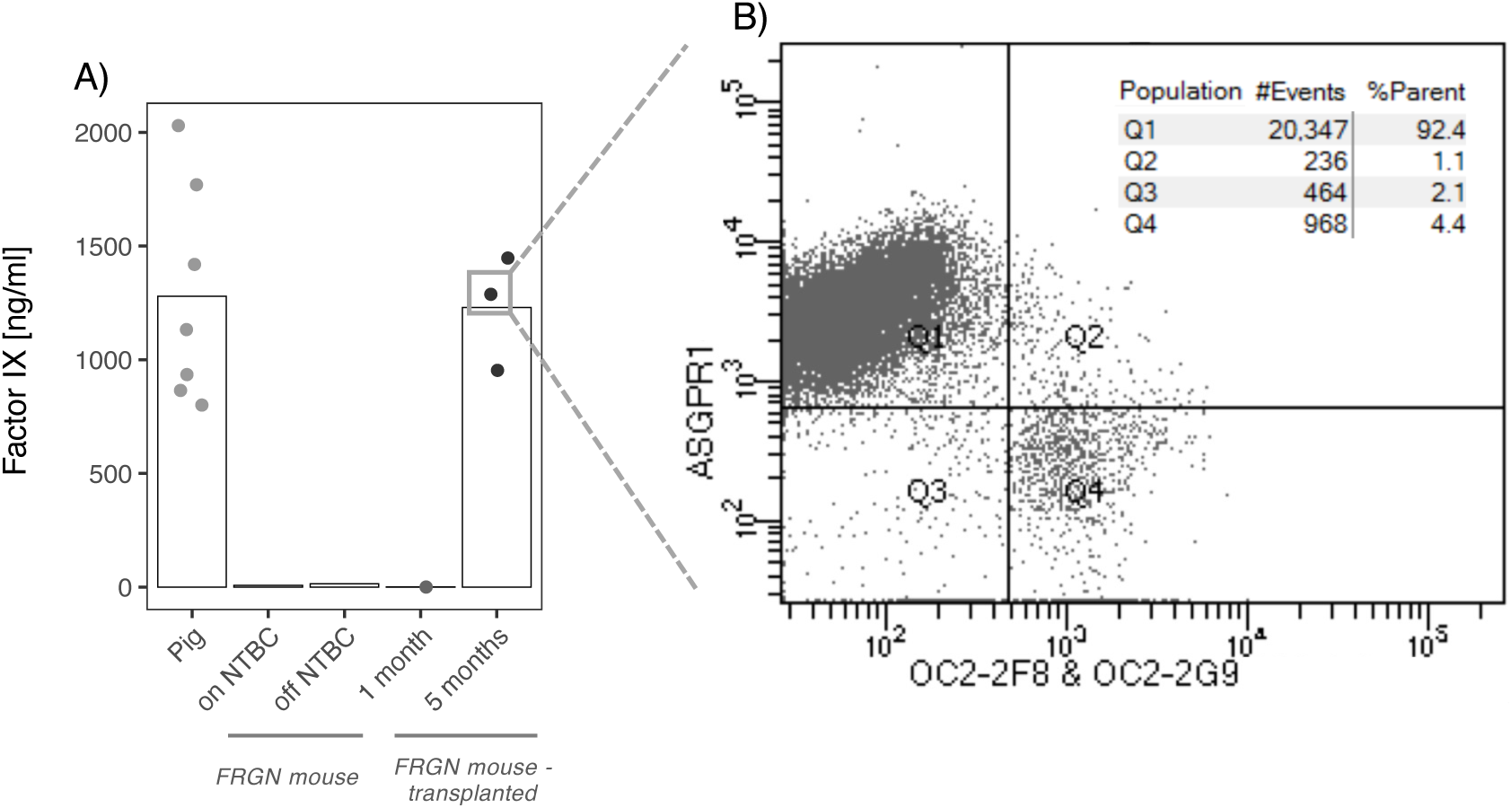
Repopulation assessment of xenotransplanted Fah-/-, Rag2-/-, Il2rg-/-, NOD (FRGN) mice (n = 3). B) Human Factor IX (hFIX) ELISA Kit detects porcine FIX in pig and pig-xenotransplanted FRGN mouse plasma and thus was used to follow repopulation. Neither murine FIX nor NTBC cycling in FRGN mice interfered with the hFIX ELISA Kit. **B**) FACS analysis (before MACS enrichment) of hepatocytes from xenotransplanted FRGN mice were harvested 5.5 months post transplantation and 2 weeks post rAAV serotype library injection. All xenograft FRGN mice were >90% repopulated with porcine hepatocytes (Q1 population). Anti-ASGPR1 antibodies were used to detect porcine hepatocytes; anti-OC2-2F8 and anti-OC2-2G9 to detect murine hepatocytes.

### *In vivo* screening using a DNA-barcoded AAV library

Highly repopulated mice (n = 3) were administered 2 x 10^13^ vector genomes (vg)/kg of the DNA-barcoded rAAV-library via tail vein (TV) injection. The rAAV-library contains 47 AAV (see Supplementary Table S3) capsids including the common AAV serotypes and genetically engineered capsids that potentially exhibit enhanced transduction in several organs including the liver, central nervous system and eyes. In this approach, AAV capsids package the AAV-CAG-BC vector genomes whose Virus Bar Codes (VBCs) are unique to each capsid. The AAV-CAG-BC genomes carry the ubiquitous CAG promoter that drives barcode expression as mRNA. This design allows the assessment of relative transduction efficiency at both DNA and RNA levels. (Figure 3A). After an incubation period of two weeks, hepatocytes were harvested and depleted of residual mouse hepatocytes by MACS. DNA and RNA were extracted from the MACS-purified porcine hepatocytes and vector genome VBCs and their transcripts were PCR-amplified by DNA-PCR and reverse transcription-PCR (RT-PCR), respectively. The resulting PCR products were then subjected to NGS-based AAV Barcode-Seq analysis as described elsewhere^44, 46^. The AAV Barcode-Seq analysis could determine AAV vector genome copy number and transduction efficiency achieved by each AAV capsid in the library relative to those of AAV9, the benchmark capsid. We used AAV9 as the benchmark capsid due to its robust ability to transduce various organs following systemic delivery^44^ (Figure 3B). AAVrh20, AAVLK03, AAV3, and AAVsh10 showed a greater than 2-fold change (2.3±0.1-, 2.2±0.2-, 2.2±0.1-, and 2.1±0.7-fold change, respectively) of DNA quantity relative to AAV9, whereas only AAVrh20, AAVLK03, and AAVAnc80 have a greater than or equal to 1.5-fold change (2.1±0.3-,1.7±0.4-, and 1.5±0.5-fold change, respectively) of RNA quantity. Therefore, with this high throughput *in vivo* method we identified AAVrh20 and AAVLK03 as the most efficient rAAV serotypes to transduce and express trans genes in porcine hepatocytes.

**Figure 3:**
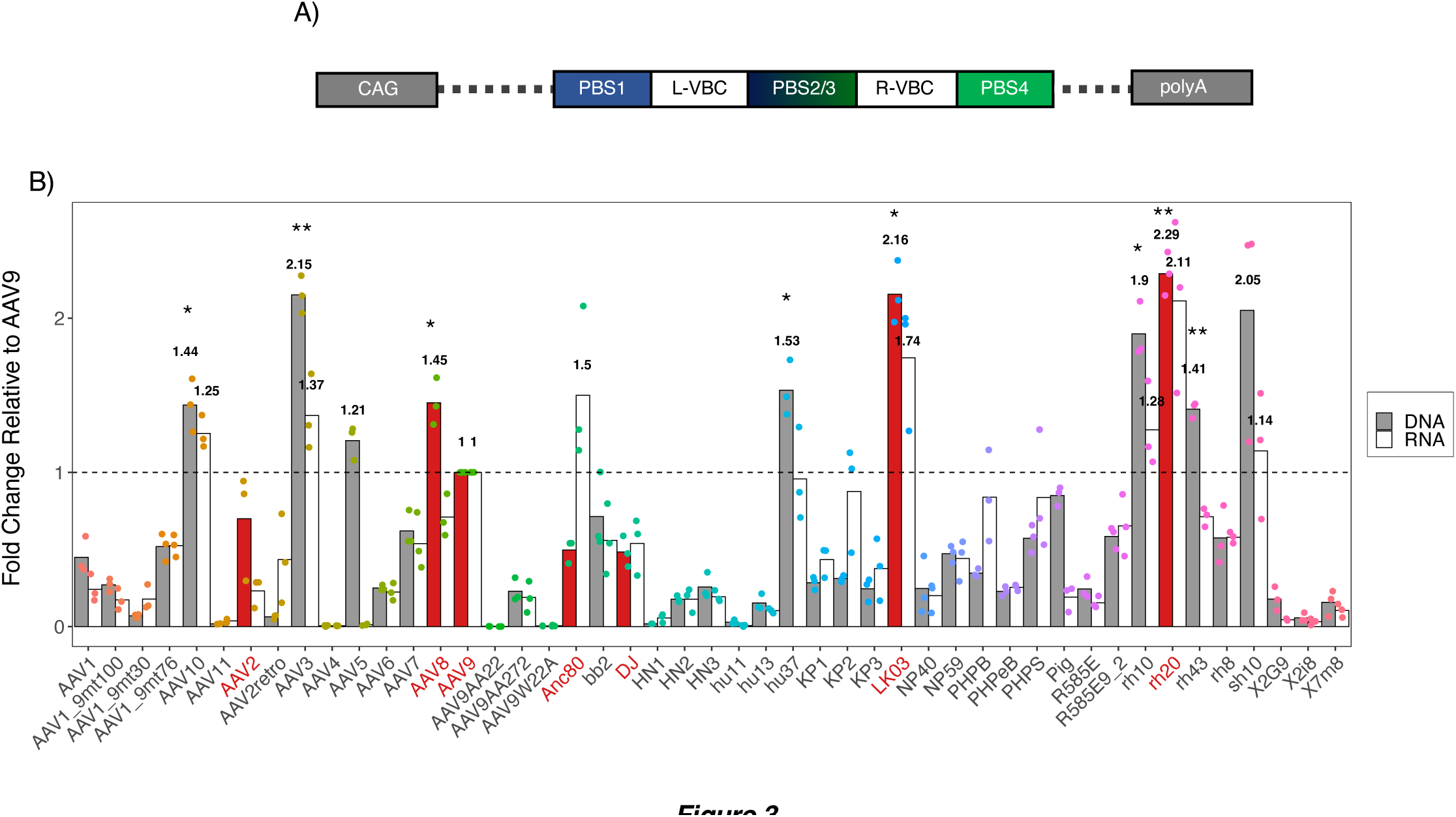
Relative gene delivery and transduction efficiency of each AAV capsid determined by AAV Barcode-Seq. **A**) Schematic of the self-complementary AAV vector encoding primer binding sites (PBS, 1-4) and viral bar codes (VBS, left and right). The vector plasmid also contain a human CAG promoter-driven nonfunctional noncoding RNA expression cassette, derived from the bacterial *lacZ* gene, with an incorporated SV40 polyadenylation signal between the two AAV2 inverted terminal repeats^46, 74^. **B**) Fah-/-, Rag2-/-, Il2rg-/-, NOD (FRGN) mice (n = 3), repopulated with porcine primary hepatocytes, were used as a model to find the most efficient AAV capsid for the porcine liver. “DNA” represents relative quantities of AAV vector genome DNA delivered by each capsid and “RNA ” represents relative transduction efficiencies at vector transcript levels. AAV9 serves as reference control, indicated by the dashed line. Statistics: paired T-test compared to PKU, ∗ p <= 0.05, ∗∗ p <= 0.01

### *In vitro* comparison of transduction efficiency of various rAAV capsids in primary porcine hepatocytes

Based on Illumina sequencing results, AAVrh20, AAVLK03, and AAVAnc80 were selected for further *in vitro* investigation, where AAV2, AAV8, AAV9, and AAVDJ were also included based on their past relevance as liver gene therapy vectors for pigs and humans and their importance as controls^19–26, 36, 37, 41^. For the total of seven selected AAV capsids, AAV2, AAV8, AAV9, AAVAnc80, AAVDJ, AAVLK03, and AAVrh20 (Supplementary Figures S1 and S3), self-complementary AAV (scAAV) vectors expressing the red fluorescent protein, tagRFP, under the control of the CMV promoter were produced and used (Figure 4A). Transduction efficiencies with these seven AAV vectors were assessed in freshly isolated 1°PH at an MOI of 20,000. These PH cells were from two 3 to 4-week-old donor pigs of both sexes and from the same litter. As depicted in Figure 4B-C, transduction levels were compared both 48 hours and seven days post-transduction based on FACS analysis and quantification of vector genome DNA copy numbers (by DNA-qPCR) and vector genome transcripts (by RT-qPCR). When comparing the relative DNA copy numbers of tagRFP of the selected AAV capsids, AAV2 and AAVDJ showed 6.3±0.8- and 1.8±0.6-fold higher relative copy numbers 48 hours post-injection and 7±5.2- and 1.6±1.1-times higher relative copy numbers seven days-post injection, respectively (Figure 4B). Thus, AAV2 and AAVDJ capsids have a higher efficiency of viral genome delivery than AAV9. On the other hand, relative percentage of tagRFP-positive cells determined by FACS analysis (Figure 4C) showed that AAV8, AAVAnc80, and AAVrh20 also had the same or higher transfection efficiency than AAV9. While the relative percentage decreased for AAVAnc80, AAVLK03, and AAVrh20 from 1.4±0.2-, 1±0.2-, and 1.1±0.3-fold to 1.1±0.2-, <1±0.3-, and 1±0.2-fold, respectively) the relative percentage for AAV2, AAV8, and AAVDJ increased (form 1.2±0.1-, 1±0.1-, and 1.2±0.1-fold to 1.4±0.05-, 1.1±0.3-, and 1.4±0.1-fold, respectively) from 48 hours to seven days post-injection. Ultimately, the efficiency to express a transgene is as important as the delivery of a vector. The relative median fluorescent intensity (MFI) of tagRFP measured by FACS shows an increase for AAV2 and AAVDJ (from 1.1±0.2- and 1.3±0.3- to 1.7±0.7- and 1.4±0.4- fold, respectively) while it decreased for AAVAnc80 (from 1.6±0.3- to 1±0.3-fold; Figure 4C). The mean MFI thus reflects the finding that AAV2, AAVAnc80, and AAVDJ not only have favorable efficiencies of delivery but also have efficient transgene expression relative to AAV9 over a period of seven days. Relative RNA copy numbers on the contrary suggest that 48 hours post-injection all selected serotypes have higher transgene expression compared to AAV9 (Figure 4B). Most significantly, 48 hours post-infection AAVDJ expresses RNA at a level 10 times higher than AAV9 and drops to 1.5 times seven days post-infection. Whereas at seven days post-infection, RNA levels of AAV2 and AAVAnc80 increase from 2.3 and 1.4 to 6.5 and >10-fold relative expression compared to AAV9, respectively. Thus, both methods, nucleic acid quantification and FACS, suggest AAV2, AAVAnc80, and AAVDJ to be the most efficient AAV capsids to transduce 1°PH *in vitro*.

**Figure 4:**
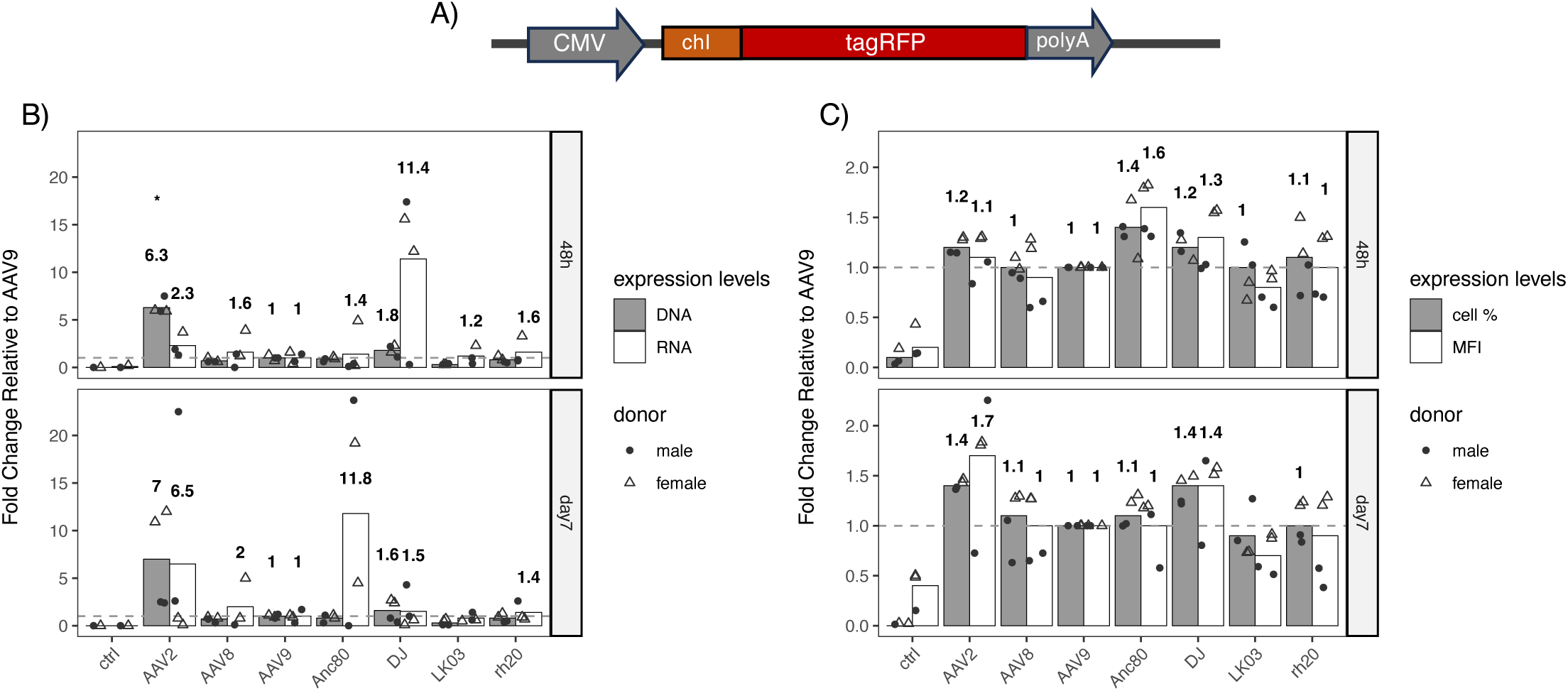
Relative transduction and expression efficiency of seven selected AAV capsids in primary porcine hepatocytes (donors = 2) transduced post-isolation (MOI of 20,000) and cultured in vitro for either 48 hours or 7 days. **A**) Self-complementary (sc) AAV vector DNA used for *in vitro* evaluation encoding tagRFP with a chimeric intron (chI) and a bovine growth hormone (bGH) polyadenylation signal under the control of the CMV promoter. The whole gene construct was flanked by scAAV2 ITRs. **B**) Relative amounts of AAV vector genome DNA and vector transcripts in AAV vector-treated primary porcine hepatocytes determined by qPCR and RT-qPCR, respectively. **C**) Relative transduction efficiencies determined by percentage of tagRFP-positive positive cells and median Fluorescent Intensity (MFI) in AAV vector-treated primary porcine hepatocytes. Fluorescence-activated cell sorting (FACS) was used to determine transduction efficiencies. AAV9 serves as the reference control, indicated by the dashed line.

Furthermore, the different methods show the benefits of single-cell evaluation using FACS since efficient viral genome delivery may not directly correlate with high transgene expression, as seen with AAVAnc80 and AAVDJ. *In vitro* testing using 1°PH, also provides insight into the timeline of intracellular processing and expression efficiency, where AAVDJ and AAVAnc80 show peaked transgene expression 48 hours and seven days post-infection, respectively, as well as differences between donors. Interestingly, when comparing absolute quantities of DNA, RNA, percentage of tagRFP-positive cells, and MFI (Supplementary Figure S2), FACS results and RNA copy numbers show noticeable differences in transgene expression between 1°PH donors for all rAAV serotypes at 48 hours post-transduction, whereas DNA copy numbers are consistent over time and between donors. Even though tagRFP was initially expressed up to 100 times higher in 1°PH from the male donor compared to the female donor, the absolute RNA copy numbers equalize seven days post-infection suggesting a transient boost in transgene expression in males immediately after infection.

## Discussion

Various AAV capsids are used in clinical trials today; nevertheless, new and improved capsids are continuously developed to further decrease viral vector and transgene immunotoxicity, increase transduction efficiency, and achieve persistent transgene expression^48–50^. To develop and test improved AAV capsids, suitable animal models are essential but also pose species dependent challenges. Besides cell entry and intracellular processing, immune responses against AAV capsids also vary greatly between species and tissues^51^. Thus, we believe the pig is a more suitable proof-of-principle large animal model for developing and testing future rAAV-based gene therapies compared to widely used rodent or canine models. Moreover, they are more accessible than non-human primates. Specifically for the liver, previous studies have utilized xenograft *FRG* mice as a system to develop, screen, and test novel AAV capsids^41, 45, 52, 53^. In this study, we established the first step toward finding a suitable AAV capsid for porcine hepatocytes and generated a chimeric *FRGN* mouse with a porcine liver repopulated to above 90%. By using a DNA-barcoded AAV capsid library, we found multiple efficient capsids for porcine hepatocytes (AAV3, AAV8, AAV10, AAVAnc80, AAVhu37, AAVLK03, AAVrh10, AAVrh20, AAVrh43, and AAVsh10; Figure 3B), with AAVLK03 and AAVrh20 capsids being the most promising gene delivery vehicles. Given that AAVLK03 efficiently transduces human hepatocytes but not mouse hepatocytes^54, 55^, our observations reaffirm that pigs, with their significant homology, serve as suitable large animal models for humans. Furthermore, the fact that AAVrh20 was originally isolated from rhesus monkey livers suggests high similarities of pigs not only with humans but also with non-human primates. The close relationship among these species is further affirmed by the affinity of multiple AAV capsids originating from rhesus monkey livers towards porcine hepatocytes, including AAV8-10, AAVrh10, AAVrh20, and AAVrh43 as seen in Figure 3. Even though the barcoded AAV library is an elegant and powerful tool, it should be kept in consideration that there may be competitive receptor interactions between the different capsids skewing the results. Ideally, the rAAV serotypes AAVLK03 and AAVrh20 should be tested *in vivo* in pigs alongside controls to confirm their superior efficiency and to determine the optimal viral dosage, immunotoxicity, and long-term impact. Testing in adult pigs would require significant resources, whereas using young piglets may be beneficial to circumvent challenges such as acquired immunity against certain AAV serotypes and requirement of large quantities of vectors. Unfortunately, using piglets to test liver-directed rAAV gene therapy has been shown to be challenging due to AAV being expressed episomally and the rapid growth of the liver^56^. Although an in-depth investigation using pigs was beyond the scope of this project, the group of Shao *et al.* demonstrated that chimeric mice can be a valuable tool to determine suitable, species-specific AAV capsids by direct comparison of hemophilia B dogs with xenograft *FRG* mice^42^. The insight gained from testing AAV capsids in 1°PH and xenotransplanted mice shown here provide initial insights and steps towards using and establishing pigs as reliable large animal models for preclinical studies.

Before and alongside *in vivo* models, immortalized cell lines have been used for decades to represent cellular mechanisms. However, due to the liver’s zonation and highly specialized functions, immortalized hepatocytes are not able to confidently replicate the liver. Thus, primary hepatocytes are considered the gold standard for *in vitro* liver-related studies, especially for metabolic processes. Recently, the group of Liu *et al.* suggested that primary hepatocytes of mice, NHPs, and humans could be used to improve and predict liver-directed rAAV gene therapies^57^. When using primary hepatocytes for *in vitro* analysis, two-step collagenase digestion is a well-established protocol. Nevertheless, it has been shown that hepatocytes undergo a pro-inflammatory response as well as mitogen-activated protein kinase (MAPK) activation, leading to a change in the hepatocyte’s phenotype upon isolation^58^, which may also affect the time required to complete post-translational modification of transgenes. Furthermore, cell surface receptors and membrane-associated proteins may be modulated by collagenase digestion as shown in rat hepatocytes^59^, which may affect viral entry into the hepatocytes. After finding suitable AAV capsids by screening a barcoded AAV capsid library, we compared AAVLK03 and AAVrh20 to other AAV capsids used for human gene therapies^2^ or as controls *in vitro* using freshly isolated 1°PH (Figure 4). Here we found AAV2, AAVAnc80 (predicted ancestor of AAV1, AAV2, AAV8, and AAV9), and AAVDJ (chimera of AAV2, AAV8, and AAV9)^37^ outperformed LK03 (contains elements of AAV1, AAV2, AAV3B, AAV4, AAV6, AAV8, and AAV9, mainly based on AAV3B) and AAVrh20. Comparing the cell entry mechanism of the capsids tested *in vitro* may provide an answer to the difference in efficiency we observed *in vitro* compared to our *in vivo* data. A majority of investigated AAV capsids use the common receptor known as the AAV receptor (AAVR, KIAA0319L)^60, 61^, which has only limited expression in the liver^62^. Whilst AAV8, AAV9^63^, presumably also AAVrh20^2^, depend on the 37/67-kDa laminin receptor (LamR) for cell entry, AAV2 and AAVDJ predominantly depend on heparan sulfate proteoglycan (HSPG)^37, 64^ but also take advantage of other membrane receptors and co-receptors^2^. AAVLK03 on the other hand, uses the hepatocyte growth factor receptor (HGFR) for cell entry like AAV3B^41, 65^. Like LamR, HGFR is a trans-membrane protein which both could be compromised during hepatocyte isolation explaining the inferior performance of AAV8, AAV9, AAVrh20, and LK03. AAV2 and AAVDJ on the contrary depend on HSPG, which is a much smaller molecule consisting of long glycan chains and potentially less affected by the collagenase digest. The investigation of this hypothesis may yield valuable insight for future *in vitro* testing using primary hepatocytes.

In conclusion, the discrepancy between *in vitro* and *in vivo* approaches highlights the difficulty to translate the two approaches when it comes to quantitative and qualitative assessment of AAV capsids suitability. It has been shown that various AAV capsids perform vastly different *in vitro* than *in vivo*^54, 66^ and emphasizes the need of standardized assays and models to compare the transduction efficiency of different capsids, most importantly including cell type and origin, incubation time, viral genome (single-stranded or self-complementary), and transgene used. The *in vitro* assessment of AAV capsids using freshly isolated hepatocytes may give insights into the mechanism of AAV vector attachment on the cell surface, cellular entry, trafficking uncoating and vector genome transcription mechanisms^41, 67, 68^ and is valuable for short term experiments. Here we also observed a donor-dependent difference of the transgene expression within the first 48 h post-infection, up to 100 times higher RNA levels and 3 times higher median fluorescent intensity (MFI) in males compared to females, which equalize seven days post-infection (Supplementary Figure S2). Primary hepatocytes may thus provide a valuable foundation for investigating gender-associated discrepancies in liver gene therapy as previously shown *in vivo* with rAAV^69, 70^. Nevertheless, primary hepatocytes should be considered with caution when used for quantitative AAV capsid comparison. On the other hand, the chimeric mouse model combined with a DNA-barcoded AAV capsid library is a valuable tool to investigate suitable AAV capsids for gene delivery in large animal models, such as pigs. By enabling a comparison of more than 40 AAV capsids, AAVLK03 and AAVrh20 were found to be promising AAV capsids for the pig liver. Efficient rAAV delivery vectors identified using the chimeric pig-mouse model enable the advanced assessment and application for more reliable pre-clinical studies in pigs, focusing on their efficacy and immunological compatibility as well as acute and long-term therapeutic effects.

## Material and Methods

### Xenografted *FRGN* Mice

The Oregon Health & Science University Institutional Animal Care and Use Committee (Portland, OR) approved all animal housing and procedures, which were performed in accordance with the approved protocols.

*Isolation of porcine hepatocytes:* Hepatocytes were isolated from porcine livers (kindly donated post euthanasia by the VirtuOHSU Simulation and Surgical Training Center at OHSU (Oregon Health & Science University)), post euthanasia by a two-step collagenase digest^71, 72^. In summary, the outer edge of a liver lobe (∼15-20 g) was removed, flushed, and stored in hepatocyte medium (high glucose Dulbecco’s modified Eagle’s medium (DMEM) substituted with 10% fetal bovine serum (FBS), 10 mM HEPES, 1% GlutaMAX, and 1% Penicillin/Streptomycin (Pen/Strep)) at 4 °C until perfusion. Within 2-3 hours, the liver lobe was subsequently perfused via an 18-gauge catheter with Ca/Mg-free Hanks’ solution (HBSS) containing 0.5 mM EGTA and subsequently with Ca/Mg-containing HBSS with 0.1 mg/ml collagenase II (Worthington). All perfusion steps were performed using pre-warmed solutions (40 °C) and a peristaltic pump (10 ml/min flow rate, for 5 and 25 minutes, respectively), ensuring a constant flow rate. Afterwards, the liver was returned to 4 °C hepatocyte medium, gently disrupted, filtered through a 100 µm and 70 µm nylon cell strainer, and washed three times at 100 *x g* for 5 minutes to generate a single-cell, hepatocyte suspension. The cell yield was estimated using 0.04% Trypan Blue, where the viability ideally exceed 80%. If the cell viability was below 70-80%, the fraction of viable cells was enriched using a Percoll gradient.

*Transplantation:* All mice were kept at controlled temperature, humidity, and a 12-hour dark-light cycle. Chow and 8 mg/L 2-2-nitro-4-trifluoromethylbenzoyl)-1,3-cyclo-hexanedione (NTBC; Yecuris Corp., Portland, OR)^39^ substituted drinking water was provided ad libitum. For transplantation, 5-8 weeks old *Fah-/-, Rag2-/-, Il2rg-/-,* NOD (*FRGN*) mice^39^ were anesthetized with 2-3% isoflurane vehicled by oxygen 1 L/min, on a heating pad (37 °C). The eyes were covered with Vitamin A, and the left flank was shaved and disinfected. The mice were transplanted via intra-splenic injection with 1 x 10^6^ viable, primary, porcine hepatocytes in 100 µl hepatocyte medium. After the procedure, mice received 2 doses 5 mg/kg carprofen (Rimadyl, Pfizer) analgesic 24 hours apart. For 2 months, the mice then were gently cycled on-off NTBC cycle, specifically on 8 mg/L NTCB for 4 days, 0.1 mg/L NTBC for 3 days, and off NTBC for 6 days. Thereafter, they were cycled 7 days on and 3 weeks off NTBC for 3 more months. To follow the degree of repopulation, the porcine factor IX (FIX) levels were measured by collecting 10 μl of whole blood in 2 weeks intervals, from the left vena saphena. After 10x dilution porcine FIX concentrations were measured using the Human Factor IX ELISA Kit (Stago) following the manufacturer’s instructions.

### DNA-barcoded AAV capsid library

*Library production:* The AAV capsid library was produced as previously described in Adachi *et al.*^44, 46^ AAV vectors were produced in HEK 293 cells using an adenovirus-free plasmid transfection method. To this end, we used pdsAAV-CAG-VBCx plasmids (where VBC is viral barcode and x is an integer identification number indicating each different viral barcode contained in each plasmid). The pdsAAV-CAG-VBCx plasmids are double-stranded AAV vector plasmids and same as pdsAAV-U6-VBCx described previously^73^ except that human U6 small nuclear RNA promoter has been replaced by the CAG promoter and SV40 polyadenylation signal has been incorporated^74^. For the benchmark AAV9 capsids, we utilized 15 VBC clones, while for other AAV capsids, we employed 2 VBC clones per capsid. Therefore, to produce the AAV capsid library, we produced a total of 107 AAV vector preparations separately. This represents 2 clones x 46 AAV capsids excluding AAV9, and 15 clones of AAV9. The individual vector titers of the 107 AAV vector preparations before purification were determined by a quantitative dot blot assay^74^. Subsequently, the 107 AAV vector preparations were mixed at approximately the same ratio and subjected to two rounds of cesium chloride ultracentrifugation followed by dialysis against phosphate-buffered saline. The final AAV library titer was determined by a quantitative dot blot assay.

*Library administration:* Throughout NTBC cycling, porcine FIX levels were determined occasionally to monitor the degree of repopulation. Once the porcine FIX levels of transplanted mice stayed > 1000 ng/mL for two consecutive measurements, mice (n = 3) were put on a low (0.7%) phenylalanine diet without NTBC. After two weeks, each mouse received 2 x 10^13^ vg/kg (lot AAV752C, Nakai Lab) of DNA-barcoded AAV-serotype library in 200 µl, via tail vein injection. Mice were incubated with AAVs for 2 weeks, after which hepatocytes were harvested by two-step collagenase digestion (described above) using a 4 mL/min flow rate. Harvested hepatocytes were then enriched for porcine hepatocytes prior to DNA and RNA extraction. Using the MACS Mouse Cell Depletion Kit (Miltenyi Biotec), cells were labelled according to manufacturer’s instructions and sorted by depletion of murine hepatocytes using the autoMACS Pro Separator. All fractions, unsorted, sorted, and depleted fractions, were labelled with mouse anti-ASGPR1-PE (1:20 dilution, to detect porcine hepatocytes) (BD Pharmigen), rat anti-OC2-2F8, and rat anti-OC2-2G9 primary antibodies (both 1:10 dilution, to detect murine hepatocytes) and mouse anti-rat Alexa-647 secondary antibody (1:200 dilution) (Jackson ImmunoResearch) to be analyzed by FACS to validate the degree of repopulation and successful enrichment of porcine hepatocytes (>95%).

*AAV Barcode-Seq:* For Illumina sequencing, total DNA and total RNA were extracted from one million cells of the porcine hepatocyte-enriched fraction using MasterPure DNA Purification Kit (Epicentre) and RNAzol RT (Molecular Research Center, Inc.) according to the manufacturer’s instructions, respectively. RNA was then treated with TURBO DNase (Invitrogen) for 1.5 hours at 37°C, and reverse transcribed into cDNA using High-Capacity cDNA Reverse Transcription Kit (ThermoFisher), also according to the manufacturer’s instructions. AAV vector genome VBCs and their transcripts, with and without reverse transcriptase treated samples, were then amplified by PCR using Platinum DuperFi DNA Polymerase (Invitrogen) and sample-specific barcode (SBC) primer sets (denaturation: 10 seconds at 98°C, annealing: 15 seconds at 60°C, elongation: 30 seconds at 72°C). Ultimately, the PCR products were subjected to NGS-based AAV Barcode-Seq analysis^44, 46^. Relative AAV vector genome DNA copy numbers in cells delivered by each AAV capsid were determined by AAV DNA Barcode-Seq as previously described^44^. Relative transduction efficiencies mediated by each AAV capsid-derived vector, measured at vector genome transcript levels, were determined by AAV RNA Barcode-Seq as previously described^46^. For RNA Barcode-Seq, the barcode sequence-dependent differences in Illumina sequencing readouts were considered. In brief, we made two AAV9-CAG-VBCx libraries that contain all the AAV-CAG-VBCx genomes packaged with the same AAV9 capsid. These two libraries were primarily the same but produced independently from two independent pools of all the pdsAAV-CAG-VBCx plasmids mixed at an equimolar ratio. Each of the two AAV9-CAG-VBCx virus libraries was intravenously injected into 8-week-old C57BL/6J male mice (n=3 each) to obtain an independent, duplicated set of data each of which was obtained from 3 mice. The livers were harvested from the library-injected mice 6 weeks post-injection. Liver RNA was extracted and VBC RNA barcode transcripts were amplified by RT-PCR. The RT-PCR products were then subjected to the Barcode-Seq analysis, which provides output RNA barcode reads. The AAV-CAG-VBCx vector genomes were extracted from each of the two AAV9-CAG-VBCx virus library stocks and the VBC DNA barcodes were amplified by DNA-PCR, which provides input DNA barcode reads. The correction factors obtained by the ratio of output RNA barcode reads and input DNA barcode reads were used to cancel out the barcode sequence-dependent differences in the RNA barcode read count data.

### *In vitro* assay using primary porcine hepatocytes

*Generation of AAVs:* Capsid sequences encoded by pAAV2/2, pAAV2/8, pAAV2/9n, and pAnc80L65AAP (catalog 104963, 112864, 112865, 92307, Addgene), as well as pAAV-DJ and pAAV-LK03 (gift from Mark Kay lab, Stanford University), pAAV-rh20 (purchased from the UPenn Core Facility), were used to package a self-complementary CMV-RFP vector. The self-complementary CMV promoter-driven tagRFP expression vector was constructed using *AgeI* and *XbaI* restriction of pscAAV-2-hCMV-chI-mRuby3-SV40p(A) (provided by Jean-Charles Paterna, University of Zurich Viral Vector Facility (VVF)) and subsequent InFusion cloning of tagRFP. The protocol for the AAV production was adapted from Addgene and performed with the guidance and help of University of Zurich VVF. Briefly, for each AAV capsid vector, ten 15 cm tissue culture dishes of HEK293T cells were triple-transfected at a molar ratio of 1:1:1. Cells and supernatant were harvested 5-6 days post-transfection, cell pellets were lysed and Benzonase-treated, and all supernatants were polyethylene glycol (PEG)-precipitated overnight. Finally, AAV was ultracentrifuged using an iodixanol gradient, concentrated and buffer exchanged with a 100 kDa molecular weight cut-off filter, and quantified by Qubit^75^.

*In vitro transduction:* Hepatocytes for *in vitro* experiments were isolated from 3 to 4-week-old piglets (males and females) according to the two-step collagen digestion protocol described above. The State Veterinary Office of Zurich approved all animal experiments. The animals were euthanized according to the guidelines of the Swiss Law of Animal Protection (licenses ZH108-2019), the Swiss Federal Act on Animal Protection (1978), and the Swiss Animal Protection Ordinance (1981). Primary porcine hepatocytes were diluted to a concentration of 1 x 10^6^ cells/mL in hepatocyte media (described above), transduced with a Multiplicity of Infection (MOI) of 20,000, and 2 mL/well cell-virus suspension was seeded into 50 µg/mL rat tail collagen type I (Sigma Aldrich, St. Louise, MO USA) coated 6-well plates. After AAV vector infection, the medium was replaced with Hepatocyte Culture Medium (HCM; Lonza, Walkersville, MD USA), which was changed every 24 hours up to the 48 hours and 7 days time point. Transduction assays of each AAV capsid was performed in duplicates for both hepatocyte donors, and each duplicate was split into three aliquots for DNA, RNA and FACS quantification. DNA and RNA aliquots were snap-frozen in liquid nitrogen and stored at -80°C until nucleic acid extraction. Replicates for FACS measurement were fixed in 1% paraformaldehyde (PFA) until quantification.

*FACS analysis:* Transduction and expression efficiency were measured by RFP detection using FACS Canto II (BD) and all data was analyzed with FlowJo software.

*qPCR and RT-qPCR analysis:* Total DNA and total RNA were extracted according to the manufacturer’s recommendation using DNeasy Blood & Tissue Kit (Qiagen, Hilden, Germany) and RNAzol RT (Molecular Research Center Inc., Cincinnati, OH USA), respectively. The nucleic acid concentrations were measured using Nanodrop, 100 ng gDNA was used for qPCR, and 1.2 µg RNA was used to produce cDNA. RNA was then treated with TURBO DNase (Invitrogen) for 1.5 hours at 37°C, and reverse transcribed into cDNA using High-Capacity cDNA Reverse Transcription Kit (ThermoFisher), also according to the manufacturer’s instructions. Total DNA and cDNA, with and without reverse transcriptase treated samples, were then amplified by qPCR using TaqMan Universal PCR Master Mix (Applied Biosystems) and tagRFP primer set (Frd: 5’- TCTGCAACTTCAAGACCACATA-3’, Rev: 5’-TCGGCCTCCTTGATTCTTTC-3’, probe: 5’-FAM- ACCCGCTAAGAACCTCAAGATGCC-TAMRA-3’). For qPCR measurement, we used ABI PRISM 7900 sequence detector (Applied Biosystems; Stage 1: 2 minutes at 50°C, Stage 2: 10 minutes at 95°C, Stage 3 (40x Repeats): 15 seconds at 95°C and 60 seconds at 60°C). SDS 2.4 (Applied Biosystems), and for analysis Sequence Detection System (Life Technologies).

## Data Availability Statement

Additional data from this study are available on request from the first and/or corresponding authors (MW and BT).

## Acknowledgements

We thank Dr. Jean-Charles Paterna from the Viral Vector Facility of the University of Zurich for his expertise and support to produce AAV. This work was supported by the Swiss National Science Foundation (310030_162547 to B.T. and mobility grants in projects CRSII5_180257/2 to M.W.) and Public Health Service grants (U01DK123608 to M.G. and H.N.). Furthermore, this work also used the Extreme Science and Engineering Discovery Environment (XSEDE), specifically the Bridges-2 system^76^ (National Science Foundation grant ACI-1548562 and award number ACI-1928147 at the Pittsburgh Supercomputing Center (PSC))

## Author contributions

M.W., A.T., K.A., B.L., and L.W. performed experiments. M.W. H.N. and K.A. analyzed data. M.W, M.G., and B.T. conceived and managed the project. M.W., and B.T. wrote the manuscript with the contribution from all authors. All authors read and approved the manuscript. M.G. and B.T. are co-correspondent authors.

