## Supplementary material for "AAV Capsid Screening for Translational Pig Research Using a Mouse Xenograft Liver Model": Supplamentary File

| donor (gender) | FRGN mice transplanted | porcine FIX positive | % successful repopulation |
|---|---|---|---|
| *Sus domesticus*, fresh (female) | **5 /** 4 | **2** / 0 | **40%** / 0% |
| Gottingen Minipig, cp (male) | **3 /** 14 | **1**/ 2 | **33%**/ 14% |
| *Sus domesticus*, cp (female) | **14 /** - | **4** / - | **29%** / - |
| total | **22** / 18 | **7** / 2 | **31%** / 11% |

***Table S1:*** *Summary of xenotransplantation and repopulation success in FRGN mice using porcine hepatocytes (fresh or cp = cryopreserved). Hepatocytes form Gottingen Minipig were aquired commercially where hepatocytes form Sus domesticus were isolated in-house. Treatment of number of recipients:* ***bold*** *= no Ad:uPA; normal = Ad:uPA.*

| mouse ID | pre-depletion (%) | post-depletion (%) | body weight (g) | liver weight (g) |
|---|---|---|---|---|
| noID | 92.4 | 97.2 | 24.9 | 3.6 |
| 2TR | 97.1 | 99.6 | 26.2 | - |
| 2BR | 95.6 | 99.6 | 28 | 5.1 |

***Table S2:*** *FACS quantification of porcine hepatocytes after hepatocyte extraction before and after depletion of mouse cells using MACS.*

| serotype | origin |
|---|---|
| AAV1 | human/non-human primate |
| AAV2 | human |
| AAV3 | human |
| AAV4 | non-human primate |
| AAV5 | human |
| AAV6 | AAV1 and AAV2 hybrid |
| AAV7 | rhesus macaque |
| AAV8 | rhesus macaque |
| AAV9 | non-human primate |
| AAV10 | cynomolgus monkey |
| AAV11 | cynomolgus monkey |
| AAV1_9mt30 | engineered |
| AAV1_9mt76 | engineered |
| AAV1_9mt100 | engineered |
| AAV2retro | engineered |
| AAV9AA22 | engineered |
| AAV9AA272 | engineered |
| AAV9W22A | engineered |
| rh8 | rhesus macaque |
| rh10 | rhesus macaque |
| rh20 | rhesus macaque |
| rh43 | rhesus macaque |
| sh10 | engineered |
| Anc80 | engineered |
| bb2 | babboon |
| DJ | shuffled |
| HN1 | engineered |
| HN2 | engineered |
| HN3 | engineered |
| hu11 | human |
| hu13 | human |
| hu37 | human |
| KP1 | in vivo directed evolution |
| KP2 | in vivo directed evolution |
| KP3 | in vivo directed evolution |
| LK03 | in vivo directed evolution |
| NP40 | shuffled |
| NP59 | shuffled |
| PHPB | engineered |
| PHPeB | engineered |
| PHPS | engineered |
| Pig | NA |
| R585E | engineered |
| R585E9_2 | engineered |
| 2G9 | engineered |
| 2i8 | domain swapping |
| 7m8 | engineered |

***Table S3:*** *47 serotypes of the AAV-serotype library and their corresponding origin, natural or engineered*

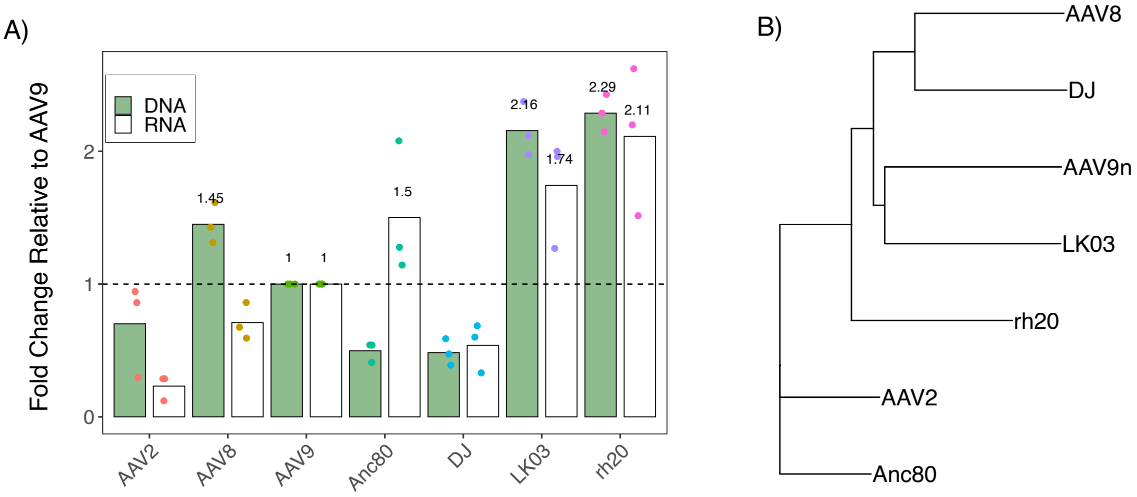


***Figure S1:*** *Selected AAV capsids for in vitro testing. A) Capsids from AAV2, AAV8, AAV9, AAVAnc80, AAVDJ, AAVLK03, and AAVrh20 were chosen based on their efficiency or as controls for further in vitro comparison using primary porcine hepatocytes. B) Phylogenetic tree representing the genetic relation of the seven chosen capsids.*

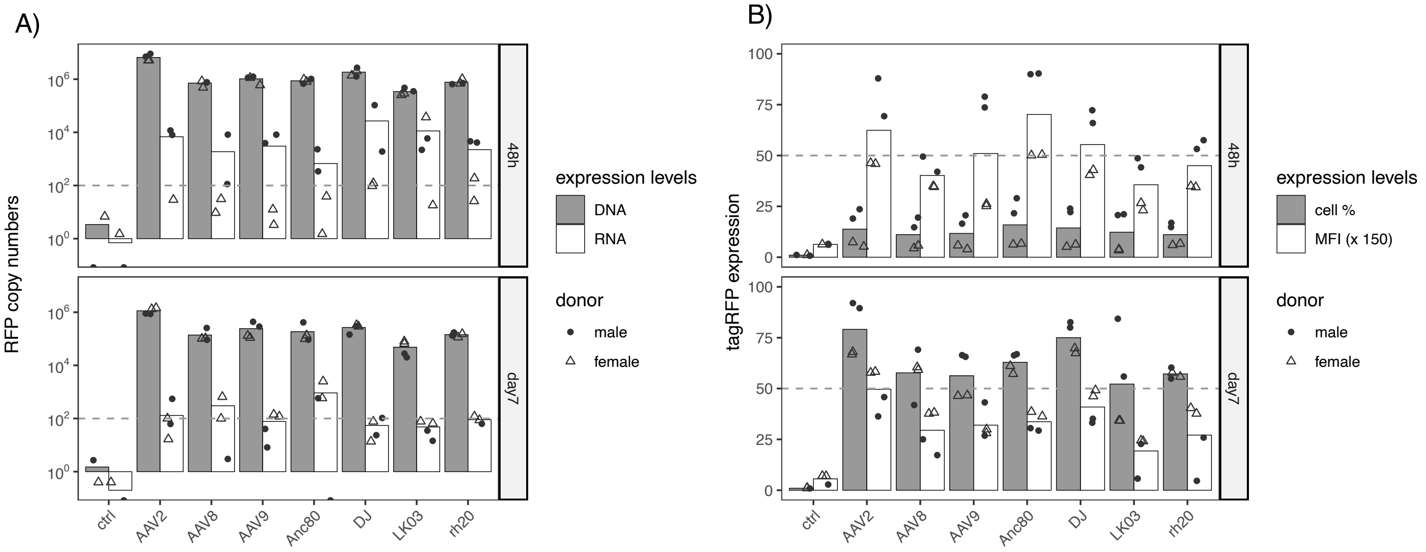


***Figure S2:*** *absolute transduction and expression efficiency of seven selected rAAV serotypes in primary porcine hepatocytes (donors = 2) transduced post-isolation (MOI 20,000 vg/cell) and cultured in vitro for 48 hours or 7 days. A) Relative mean DNA and RNA amounts post-transduction in primary porcine hepatocytes measured using qPCR and RT-qPCR, respectively. B) Relative mean of tagRFP-positive single cells and Median Fluorescent Intensity (MFI) post-transduction in primary porcine hepatocytes measured using fluorescence activated cell sorting (FACS). Dashed line represents 50% tagRFP positive cells, 7500 MFI, and* $10^2$ *tagRFP copy numbers per 100 ng DNA or 1200 ng RNA.*

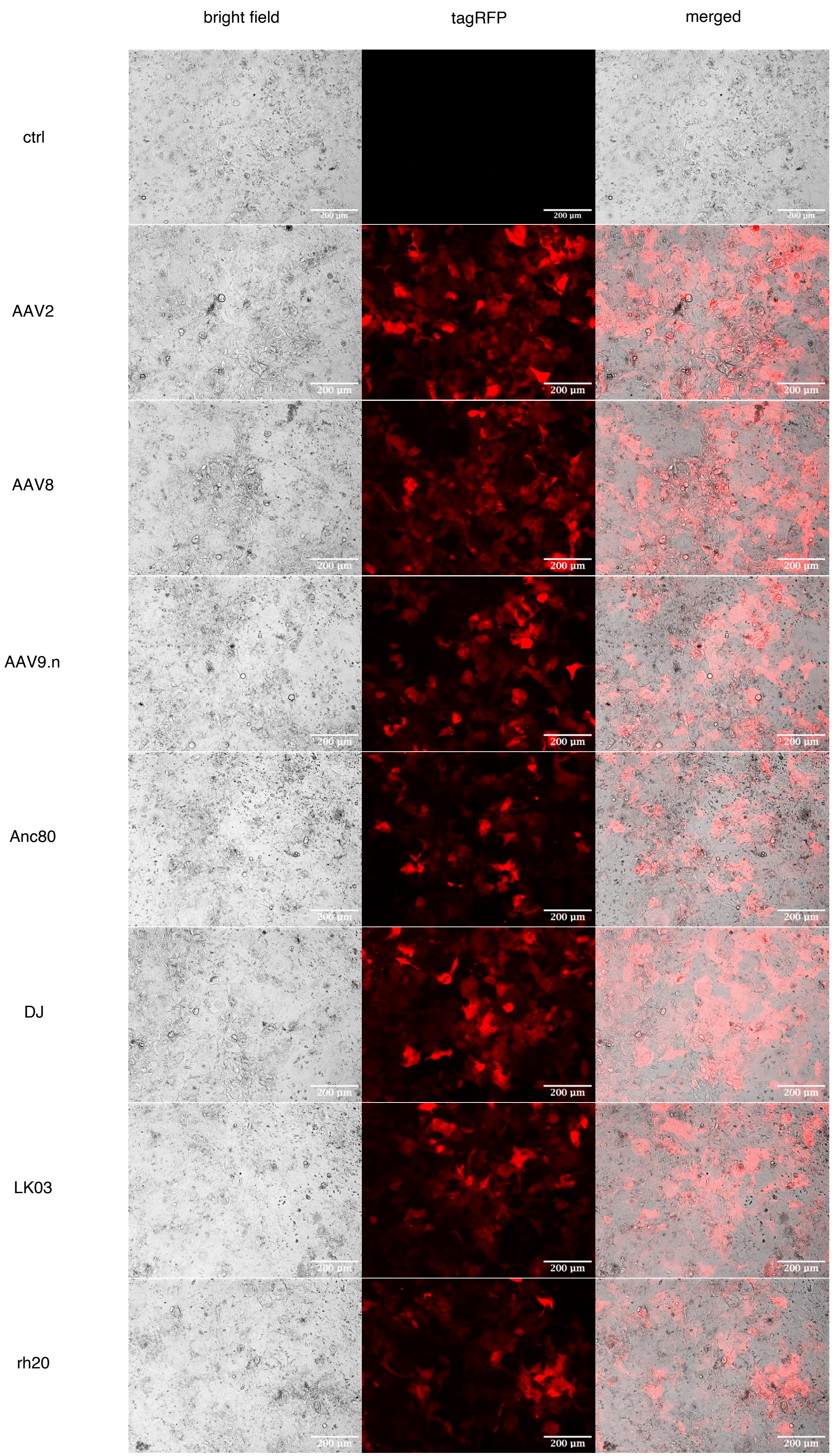


***Figure S3:*** *Visualization of tagRFP expression in primary porcine hepatocytes transduced with selected AAV-serotypes, seven days post-transduction. Scale bars, 200 µm.*
